# The effect of drug dose and timing of treatment on the emergence of drug resistance *in vivo* in a malaria model

**DOI:** 10.1101/2020.01.27.921940

**Authors:** Monica M. Acosta, Joshua T. Bram, Derek Sim, Andrew F. Read

**Author notes:** corresponding author. University of Michigan, Ann Arbor, MI, 48109, USA. LSA Complex Systems, University of Michigan, Ann Arbor, MI, 48109. Perelman School of Medicine, University of Pennsylvania, Philadelphia, PA, 19104, USA. Joshua T. Bram (e-mail). Derek Sim (e-mail). Andrew F. Read (e-mail).

## Abstract

**Background and objectives:** There is significant interest in identifying clinically effective drug treatment regimens that minimize the *de novo* evolution of antimicrobial resistance in pathogen populations. However, *in vivo* studies that vary treatment regimens and directly measure drug resistance evolution are rare. Here, we experimentally investigate the role of drug dose and treatment timing on resistance evolution in an animal model.

**Methodology:** In a series of experiments, we measured the emergence of atovaquone-resistant mutants of *Plasmodium chabaudi* in laboratory mice, as a function of dose and timing of treatment with the antimalarial drug atovaquone.

**Results:** Increasing the concentration of atovaquone increased the likelihood of high-level resistance emergence. Treating very early or late in infection reduced the risk of resistance, likely as a result of population size at time of treatment, but we were not able to exclude influence of the immune response in the latter. When we varied starting inoculum, resistance was more likely at intermediate inoculum sizes, but this did not correlate directly with population sizes at time of treatment.

**Conclusions and implications:** (i) Higher doses do not always minimize resistance emergence and can result in competitive release of parasites with high-level resistance. (ii) Altering treatment timing affects the risk of resistance emergence, but not as a simple function of population size at the time of treatment. (iii) Finding the ‘right’ dose and ‘right’ time to maximize clinical gains and limit resistance emergence can vary depending on biological context and was non-trivial even in our simplified experiments.

## INTRODUCTION

There is widespread agreement that appropriate antimicrobial use is critical for minimizing the emergence and spread of antimicrobial resistance. This belief is encapsulated in the mantra that resistance development can be inhibited by the right drug at the right time at the right dose for the right duration [1,2]. Here we focus on two of those factors – timing and dose – and ask experimentally how they affect the emergence of *de novo* drug resistance in an animal model. Our goal is to further the underlying science needed to identify treatment regimens which minimize resistance emergence while maximizing host health. Surprisingly few experimental studies have addressed the issue *in vivo* [3,4].

The *de novo* evolution of drug resistance poses a significant challenge for the management of many diseases [5,6]. In some cases, *de novo* resistance contributes to treatment failure, as with cancer [7] and several quickly evolving viral [8–10] and bacterial infections [11–13]. In other cases, *de novo* evolution is important as a source for resistance that is then transmitted and spread within a population [14]. For *de novo* resistance to emerge in a patient, two conditions must be satisfied. First, resistant variants must appear, either by mutation or, in the case of some bacteria, by horizontal gene transfer. Second, resistant sub-populations must then expand to densities within their host that trigger symptoms and/or become transmissible. Both of these processes are impacted by drug treatment.

A common belief is that high enough drug doses can prevent resistance [15–20] because higher drug concentrations are more likely to kill resistant mutants [15,21,22]. In the limit, this is clearly true, but if concentrations sufficient to kill all resistant mutants cannot be achieved, populations of surviving mutants rapidly expand to fill the niche vacated when pathogens are killed by chemotherapy [3,4,23–25]. This process of niche expansion (also called ‘competitive release’) has been demonstrated in animal disease models [24,26–28] and in humans [29,30]. Competitive release means that in the simplest case, the relationship between drug dose and resistance emergence is an ‘inverted U’: at very low doses, there is no selection for resistance, at high doses, everything is killed, and in between, resistance evolution is promoted [3,4,31]. The inverted U has been readily observed *in vitro* [4], but there has been very limited *in vivo* testing [3,4]. *In vivo* testing is important for at least three reasons. First, immunity is a key determinant of both antimicrobial efficacy and pathogen population sizes [4,32–36]. Second, within-host density-dependent population regulation is a key determinant of competition between wildtype and mutant populations [26,37]. Third, and perhaps most important, resistance-minimization strategies have to be considered in the context of patient health, which by definition can be directly measured only *in vivo*.

Another common belief is that the earlier treatment begins, the less likely resistance is to arise [21,38,39]. The thinking underlying this belief, is that adaptive evolution (here resistance) proceeds faster in larger populations because larger populations are more likely to contain mutations on which selection can act [40–43]. This is indeed well verified *in vitro*, where resistance is more likely to emerge when larger populations are treated with antimicrobials (e.g. ref. 42). However, theoretical analyses have pointed to important complexities that can arise *in vivo* [32–34,44,45]. For example, density-dependent immune components may be differentially affected by pathogen exposure, which early treatment can truncate, restricting control of resistance in the context of developing immunity [34]. Furthermore, increases in population size do not always result in increased levels of genetic diversity. For example, small differences in the growth rate between resistant mutants and wild type parasites can reduce genetic diversity over time [46]. The few *in vivo* tests of which we are aware [38,39,47–50] involved early or late drug treatment or infections initiated with small or large inocula [39,49]. In these binary cases, resistance typically emerged less readily if infections were treated early or had fewer pathogens to begin with, consistent with the idea that larger populations are more likely to contain resistant mutants.

Here, we use a rodent model of malaria to test the effects of drug dose on the *de novo* emergence of resistance and attempt to disentangle the impact of treatment timing and pathogen population size. Atovaquone is a highly effective antimalarial drug when used in combination with proguanil, but when used as a monotherapy it fails in up to 30% of patients because of resistance evolution [51,52]. Atovaquone resistance is due to easily identified point mutations in the mitochondrial *cytochrome b (cytb)* gene which confer high-level resistance [51,53], and likely arises rapidly due to the multi-copy nature of mitochondrial DNA [54] and unique mode of replication involving extensive copy correction [53,55]. We reasoned that a strong resistance phenotype that readily evolves and is conferred by easily assayed genetic changes would make possible an experimental analysis of the impact of dose and timing on the probability of resistance emergence.

## METHODOLOGY

### Experimental overview

We conducted five separate experiments (**Fig. 1, Table 1**). Two experiments explored the effects of atovaquone dose (concentration) on the probability of resistance emergence (**Fig. 1A**). In an initial experiment, we explored a set of doses and then, in a second experiment, we extended the range of doses. In two further experiments, we held atovaquone dose constant and looked at the impact of treatment timing on resistance emergence. In a final experiment, we held both dose and the timing of treatment constant and varied the number of parasites used to initiate the infections (inoculum size). This generated different parasite population sizes at the start of treatment (**Fig. 1 C-E**). In experiments 3 and 4, we varied the day post-infection of drug administration and in experiment 5, we held the day of treatment constant but varied the number of parasites used to initiate infection, so as to generate different populations sizes on the day of treatment.

**Table 1.** Experimental details. All drug treatments were performed on two successive days. In the case of experiments 3-5, 4 mg/kg was used. Mice were omitted from analysis if they died during acute stage infection or were mis-inoculated (see methods). Phenotypic resistance testing was only performed in experiments 1 and 2.

| Experimental group | Inoculum size | Treatment start | Total mice | Total in analysis | No. relapsed | No. resistance phenotyping |
| --- | --- | --- | --- | --- | --- | --- |
| ***Atovaquone dose and emergence of resistance*** |  |  |  |  |  |  |
| *Experiment 1 (C57BL/6)* |  |  |  |  |  |  |
| 0 mg/kg | 10<sup>6</sup> | 6 | 10 | 9<sup>a</sup> | - |  |
| 1 mg/kg | 10<sup>6</sup> | 6 | 10 | 10 | 10/10 | 2 |
| 2 mg/kg | 10<sup>6</sup> | 6 | 10 | 9<sup>b</sup> | 8/9 | 3 |
| 4 mg/kg | 10<sup>6</sup> | 6 | 10 | 8<sup>ab</sup> | 6/8 | 4 |
| 8 mg/kg | 10<sup>6</sup> | 6 | 10 | 9<sup>b</sup> | 5/9 | 5 |
| *Experiment 2 (Swiss Webster)* |  |  |  |  |  |  |
| 0 mg/kg | 10<sup>6</sup> | 6 | 10 | 1<sup>9a</sup> | - | 1 |
| 1 mg/kg | 10<sup>6</sup> | 6 | 10 | 8<sup>abc</sup> | 8/8 | 7 |
| 2 mg/kg | 10<sup>6</sup> | 6 | 10 | 9<sup>a</sup> | 9/9 | 5 |
| 4 mg/kg | 10<sup>6</sup> | 6 | 15 | 12<sup>2ab</sup> | 12/12 | 7 |
| 8 mg/kg | 10<sup>6</sup> | 6 | 20 | 17<sup>2acd</sup> | 15/17 | 8 |
| 12 mg/kg | 10<sup>6</sup> | 6 | 20 | 15<sup>5a2c</sup> | 15/15 | 8 |
| 18 mg/kg | 10<sup>6</sup> | 6 | 20 | 15<sup>4abc</sup> | 14/15 | 7 |
| ***Population size and timing and emergence of resistance*** |  |  |  |  |  |  |
| *Experiment 3 (Swiss Webster)* |  |  |  |  |  |  |
| 3 days PI | 10<sup>2</sup> | 3 | 7 | 7 | 0/7 | - |
| 5 days PI | 10<sup>2</sup> | 5 | 7 | 6<sup>a</sup> | 0/6 | - |
| 7 days PI | 10<sup>2</sup> | 7 | 7 | 6<sup>a</sup> | 0/6 | - |
| 9 days PI | 10<sup>2</sup> | 9 | 7 | 7 | 2/7 | - |
| 11 days PI | 10<sup>2</sup> | 11 | 7 | 6<sup>b</sup> | 6/6 | - |
| *Experiment 4 (C57BL/6)* |  |  |  |  |  |  |
| 6 days PI | 10<sup>6</sup> | 6 | 8 | 7<sup>b</sup> | 7/7 | - |
| 9 days PI | 10<sup>6</sup> | 9 | 8 | 4<sup>4a</sup> | 2/4 | - |
| 12 days PI | 10<sup>6</sup> | 12 | 8 | 6<sup>2a</sup> | 2/6 | - |
| 15 days PI | 10<sup>6</sup> | 15 | 8 | 7<sup>a</sup> | 0/7 | - |
| 18 days PI | 10<sup>6</sup> | 18 | 8 | 4<sup>3ab</sup> | 0/4 | - |
| 21 days PI | 10<sup>6</sup> | 21 | 8 | 6<sup>ab</sup> | 0/6 | - |
| *Experiment 5 (Swiss Webster)* |  |  |  |  |  |  |
| 10<sup>2</sup> inoculum | 10<sup>2</sup> | 6 | 8 | 7<sup>b</sup> | 2/7 | - |
| 10<sup>3</sup> inoculum | 10<sup>3</sup> | 6 | 8 | 5<sup>3b</sup> | 5/5 | - |
| 10<sup>4</sup> inoculum | 10<sup>4</sup> | 6 | 8 | 7<sup>b</sup> | 7/7 | - |
| 10<sup>5</sup> inoculum | 10<sup>5</sup> | 6 | 8 | 7<sup>bc</sup> | 6/7 | - |
| 10<sup>6</sup> inoculum | 10<sup>6</sup> | 6 | 8 | 7<sup>b2c</sup> | 7/7 | - |
| 10<sup>7</sup> inoculum | 10<sup>7</sup> | 6 | 8 | 3<sup>5a</sup> | 3/3 | - |

<sup>a</sup> mice which died during the acute stage of infection, <sup>b</sup> mis-inoculated mice (defined in the text). <sup>c</sup> mice which died during relapse, but are included in the analysis except where indicated. <sup>d</sup> mouse which was omitted from analysis due to unobtainable Sanger sequencing data. Number immediately preceding subscript relates to number of mice in each category.

**Fig 1.**
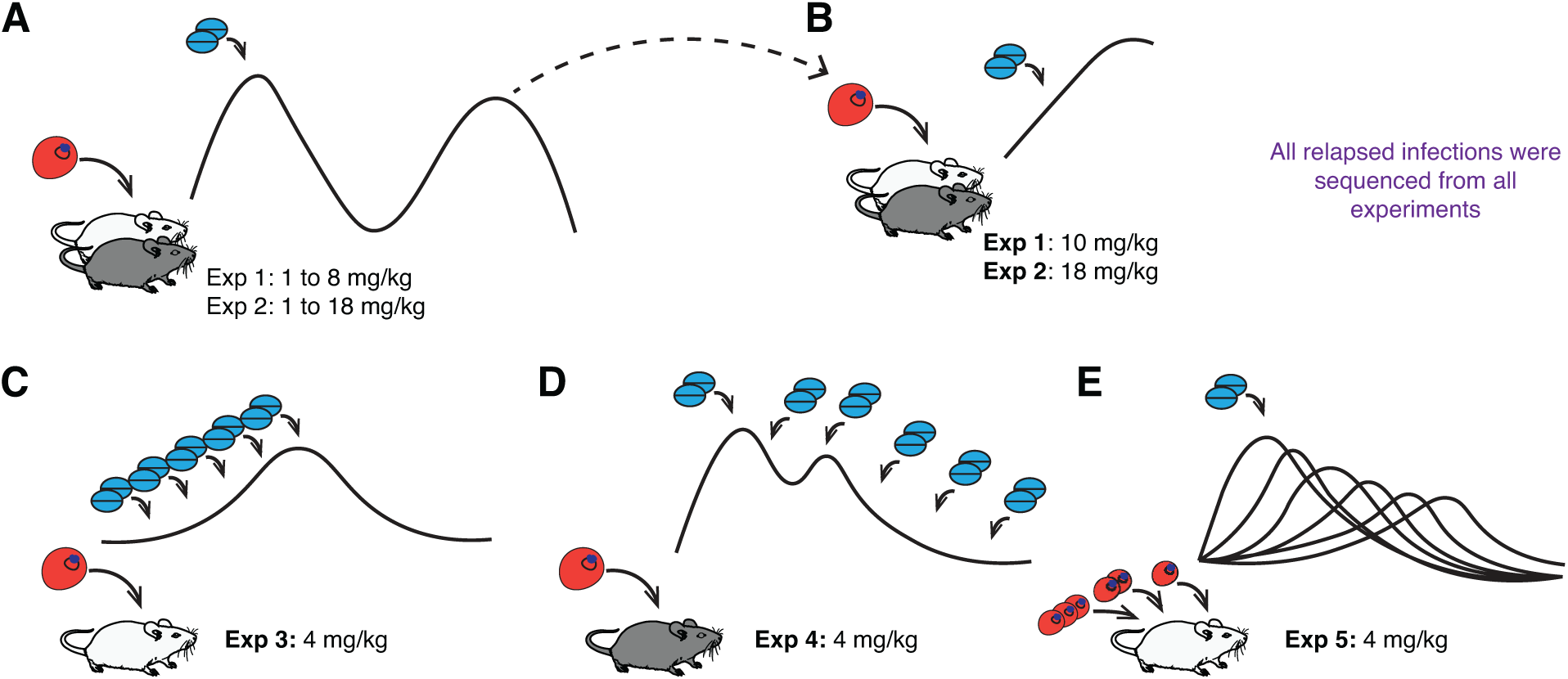
Experimental overview. **(A)** Effect of atovaquone dose and resistance emergence (experiments 1 and 2). Mice were drug treated with varying doses and monitored for relapse. A subset of relapsing infections was passaged to naïve mice **(B)** and drug-treated with a high dose to measure phenotypic resistance (see methods). All relapsed infections were sequenced at the Qo2 region of the *cytb* gene for high-level resistance. **(C-E)** Experimental manipulations of population numbers at the start of treatment. In experiment 3 **(C)** and 4 **(D)**, infections were treated at different times leading up to or following peak parasitemia, respectively. In experiment 5 **(E)**, treatment day was fixed, but different inoculum sizes used to seed infections. Mice in experiments 3-5 were monitored for relapse and similarly genotyped for high-level resistance as experiments 1 and 2. Black and white mice represent C57BL/6 and Swiss Webster standard mouse lab strains, respectively.

### General methods

#### Mice and parasite strains

All experimental mice were female, aged six to eight weeks old at the start of the experiments. Experiments 1 and 5 were conducted with inbred C57BL/6 mice, while experiments 2-4 involved outbred Swiss Webster mice (**Table 1**). Mice were housed in cages of four or five individuals and maintained on a diet of 0.05% para-aminobenzoic acid (an organic compound added to facilitate parasite growth in this model [56]) and on mouse chow (PicoLab® Rodent Diet 20). Parasites were of the AS_13p_ clone of *Plasmodium chabaudi*. This lineage of *P. chabaudi* has no prior exposure to atovaquone. All parasite inoculations were performed intraperitoneally with appropriate dilutions in citrate saline in a total volume of 100 μL.

#### Drug treatment

Atovaquone was dissolved in DMSO and up to 100 μL was inoculated intraperitoneally. All drug treatments were performed in the late morning. Control mice which did not receive atovaquone were inoculated with an equal volume of DMSO. Studies in other rodent malaria models where atovaquone resistance emerged used 5 mg/kg [51] and 14.4 mg/kg [57] as therapeutically realistic concentrations (without defining treatment durations). We chose drug doses spanning these values and chose to treat for two days. These doses are higher than the MIC reported for various other rodent malaria systems (range 0.04 mg/kg to 0.1 mg/kg) [57,58].

#### Monitoring infections

In most cases, mice were monitored daily starting three days post-infection. Exceptions were experiment 2, in which mice were sampled only on even days starting four days post-infection, and during periods late in infection in experiments 3-5, when sample frequency was changed to every two or three days. Details of blood sampling, quantification of red blood cell (via coulter counter) and parasite densities (quantitative PCR) are described elsewhere [59,60].

#### Genotyping for resistance

As genetic markers of resistance, we focused on the development of high-level resistance, which is associated with SNP mutations in the Qo2 domain of the parasite’s mitochondrially-encoded *cytb* gene [58,61–63]. As such, we amplified and Sanger sequenced a 422 base pair region encompassing the entire Qo2 region (**Supplementary Fig. S1**). Samples were analyzed and aligned to an available reference sequence of *cytb* for *P. chabaudi* strain AS (GI: 222425464) using Geneious® version 9.1.8. Lower level resistance is associated with SNPs in other domains of the *cytb* gene [57,58], but we focus on mutations in the Qo2 domain because slowing and even preventing the emergence of high-level resistance is a long-term goal of evolutionary medicine [4]. Thus, our study focuses on the impact of contrasting treatment regimens on the emergence of resistance encoded in the Qo2 domain. Further details regarding our genotypic sequencing are available in the **Supplementary Text**. A measure of phenotypic resistance was also included in our experiments in which we varied dose, which we detail below.

### Experimental details

#### Drug dose and emergence of resistance

For experiments 1 and 2 (C57BL/6 and Swiss Webster mice, respectively), hosts were infected with 10^6^ parasites and drug treated on days six and seven post-infection when parasite densities peak and mice become lethargic, anemic and lose weight. Drug doses spanned 1 mg/kg to 8 mg/kg and 1 mg/kg to 18 mg/kg for each experiment, respectively (**Table 1, Fig. 1A**). To measure the resistance phenotype, a subset of relapsed infections in all treatments (sampled haphazardly, with a goal of capturing at least three per treatment) were passaged to naïve mice (infections initiated with 10^6^ parasites, **Table 1, Fig. 1B**). These were then drug treated (10 mg/kg or 18 mg/kg, experiment 1 and 2, respectively) on days three and four post-infection and monitored until day seven post-infection. All relapsed infections were sequenced as detailed above.

#### Timing of treatment, population size and emergence of resistance

In experiments 3-5, we kept drug treatment constant at 4 mg/kg for two successive days and attempted to vary pathogen population size at the start of atovaquone treatment with three separate experimental manipulations (**Table 1, Fig. 1C-E**). In experiment 3, we infected Swiss Webster mice with 100 parasites and drug treated at various times before peak parasitemia. Inoculation with 100 parasites results in prolonged time to maximum parasitemia [64]. In experiment 4, we infected C57BL/6 mice with 10^6^ parasites, and drug treated on different days after peak parasitemia. Finally, in experiment 5, we infected Swiss Webster mice with different numbers of parasites (10^2^, 10^3^, 10^4^, 10^5^, 10^6^ or 10^7^) and drug treated on days six and seven post-infection. All relapsed infections were sequenced as detailed above, and in this case phenotypic resistance was not assayed.

### Statistics

#### Definition of parasite relapse and resistance

Relapse is the appearance of sustained parasite populations after drug treatment. We operationally defined this as parasite numbers above our assay detection threshold for greater than three consecutive sampling time points. In all cases, relapse was easily identifiable, and parasites remained above detectable levels for at least six days. Resistance was defined as relapse that resulted in any majority genotype that differed from the wildtype *cytb* sequence at the Qo2 domain. All mutations we found had been previously associated with atovaquone or were at positions in which amino acid substitutions have previously been reported to result in resistance [51,57,63]. In experiments 1 and 2 where we measured phenotypic resistance, we additionally evaluated resistance as the ability for passaged parasites to grow in the presence of atovaquone in naïve mice.

#### Analysis

All analyses and graphics were conducted in R, version 3.5.1 [65] and RStudio [66]. Parasite numbers per mouse were log10 transformed prior to all analyses. Mice that died before relapse could have occurred were excluded from all analyses. Mice for experiments 1 and 2 which were mis-inoculated with parasites, defined as those which differed over two log fold from average parasite densities on the first day of monitoring, were similarly omitted from analyses. In the case of experiments 3-5, where population size on day of treatment was the focus, we were more conservative in our definition of mis-inoculations, dropping any mice which differed over a log fold from the average parasite number on the first day of treatment (with the exception of mice which were treated on days three or six post-infection in experiment 3 which were below the level of detection at this timepoint).

Generalized linear models were used to analyze probabilities of relapse and resistance. For experiments 1 and 2, drug dose was analyzed as a continuous variable (log transformation of drug dose resulted in similar patterns, data not shown). In experiment 3, we estimated parasite numbers at the time of treatment in the earlier drug treated experimental groups where numbers were often below our qPCR detection threshold (**Supplementary Fig. S2**).

Parasite decline during drug treatment (clearance rate) and passaged parasite growth in naïve mice in experiments 1 and 2 were calculated by fitting individual linear models for each mouse (days six to ten and days three to six, respectively). In our analysis of resistance phenotypes, drug dose was log transformed to linearize relationships. We used a generalized least squares model to analyze differences in pathogen population size across experiments using the R function VarIdent assuming different variances per experiment within the gls package. Note in what follows, we restrict the word treatment to mean drug treatment and instead use the terms experimental group or manipulation to refer to differences in the timing of drug treatment or inoculum size for clarity. All p-values are for two-tailed tests.

## RESULTS

### Atovaquone dose

Our first two experiments tested the effect of atovaquone dose on resistance emergence (**Fig. 1A and B**).

Atovaquone dose altered resistance emergence in both experiments (summarized in **Supplementary Table S1, Fig. 2)**. In experiment 1 (C57BL/6 mice), treatment failure in general was prevented at higher doses, with fewer instances of relapse (χ_1,35_^2^ = 6.1, p = 0.01, **Fig. 2A**). However, resistance mutations in the Qo2 domain were as likely to emerge at any drug dose (χ_1,35_^2^ = 0.1, p = 0.7, **Fig. 2A and 3A**). By contrast, in experiment 2 (Swiss Webster mice, wider range of drug doses), treatment failure occurred in almost all cases and was unrelated to drug dose (drug dose: χ_1,75_^2^ = 0.6, p = 0.4, **Fig. 2B**), but the failures at higher doses were more likely to be due to resistance evolution (χ _1,75_^2^ = 11.5, p < 0.001, **Fig. 2B and 3A**). When we limited our analysis to doses that were common between the two experiments (1 to 8 mg/kg), we found that the impact of dose on the probability that resistant mutants emerged differed between experiments (dose*experiment: χ_3,81_^2^ = 10.16, p = 0.02).

**Fig 2.**
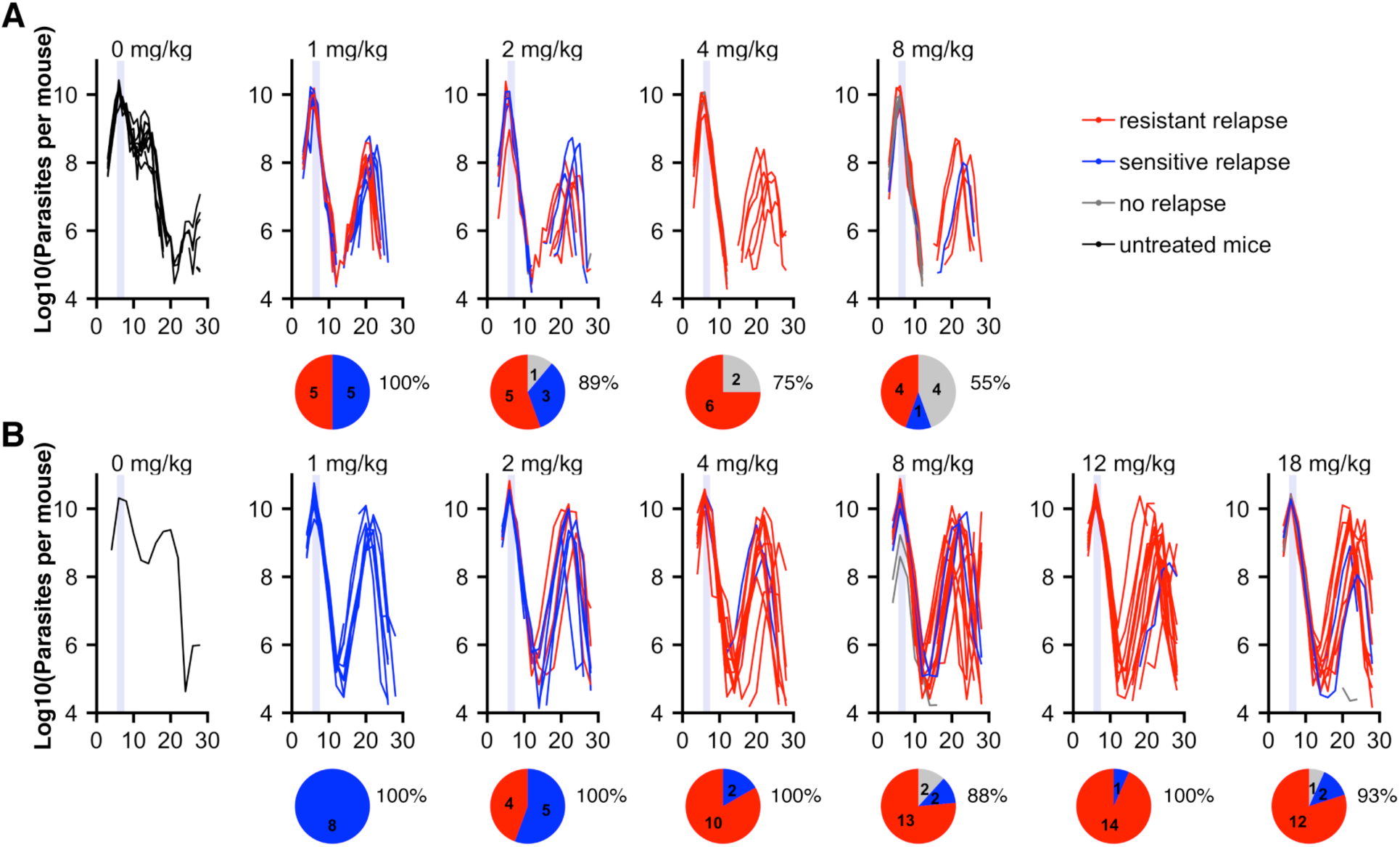
Parasite dynamics for individual mice in experiments 1-2 showing treatment failure and resistance. Parasite dynamics are shown through time for all individual mice in experiments 1 **(A)** and 2 **(B)**. Red and blue indicate the identity of relapsing parasite populations from Sanger sequencing of the Qo2 region of the *cytb* gene, with blue representing wildtype genotypes and red indicating the presence of mutation(s) known to encode high-level resistance. In grey are dynamics of non-relapsing infections. Pie diagrams show proportion and total numbers of mice in each category. Percentages show proportion of total infections which relapsed per treatment. Dynamics of surviving mice which were not drug treated are shown in black. Drug treatment timing is represented as light purple bars.

**Fig 3.**
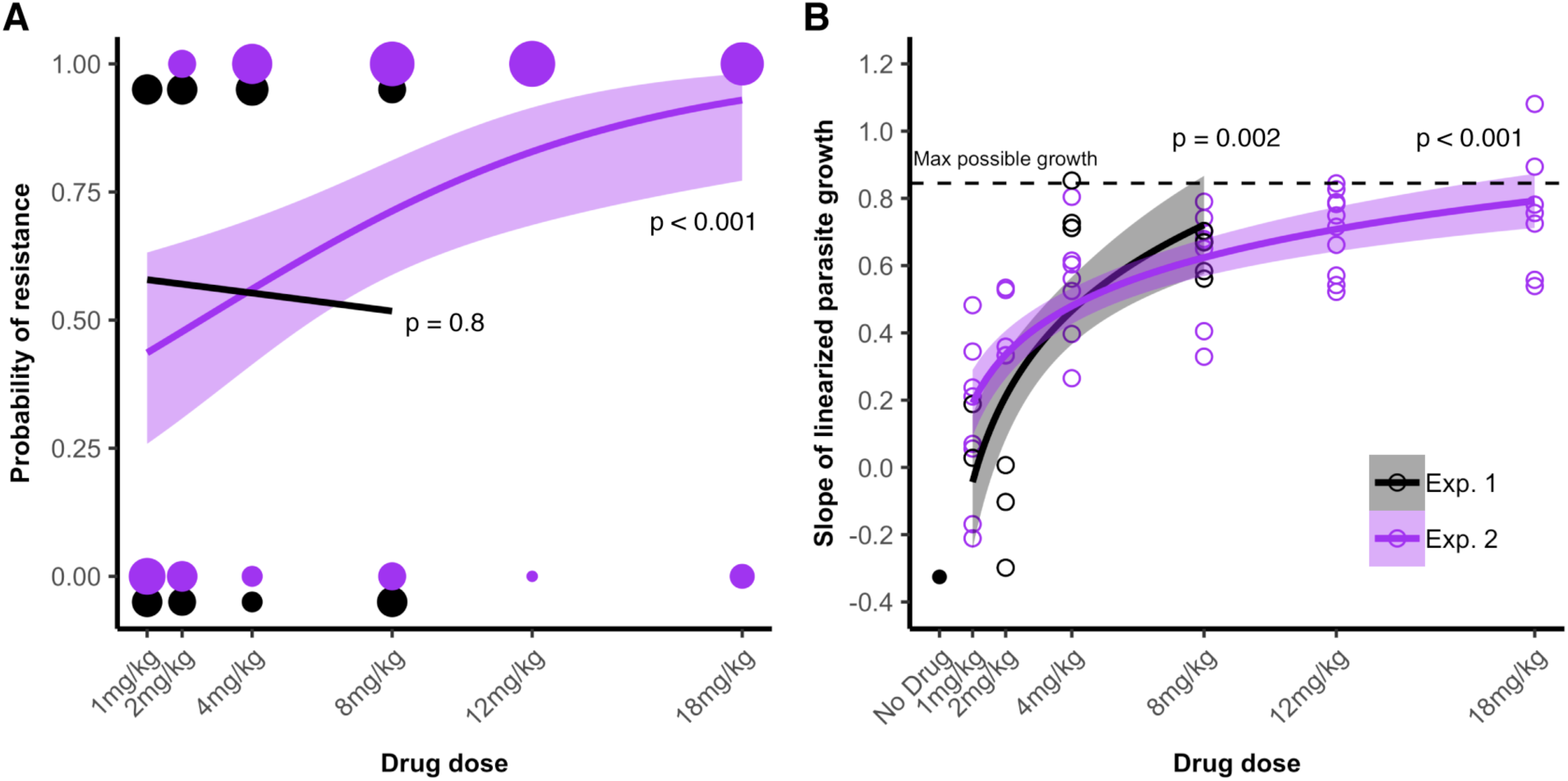
Dose-dependent emergence of atovaquone resistance. Plotted data for experiment 1 (black) and experiment 2 (purple). **(A)** Relapsing populations after drug treatment were genotyped at the Qo2 region of the *cytb* gene and classified as wildtype (0) or drug-resistant (1). Probability of resistance evolution (mice with relapsing resistant infections out of all mice in each treatment) are plotted. Size of circles correspond to number of mice. Lines are from GLM models fitted for each experiment. **(B)** A subset of relapsing infections was passaged to naïve mice and atovaquone-treated. Plotted points show population growth rates in mice. Lines represent linear models of dose (log10 transformed) against parasite growth for each experiment. The dotted line represents the growth rate calculated for a burst size of seven merozoites per infected red blood cell (representing an average burst size for *P. chabaudi*). Error widths for lines shown are 95% confidence intervals.

To correlate our initial genotypic measure of resistance with phenotypic drug susceptibility of relapsing infections, a subset of infections (see methods) were passaged to naïve mice and treated with a high dose of atovaquone for two days early in infection. A total of 14 (48% of total relapses) for experiment 1, and 59 infections (77% of total relapses) for experiment 2 were tested in this manner. Parasites that had been genotyped as resistant grew better in the presence of the drug than those genotyped as wildtype (F_1,57_ = 50.5, p < 0.001). Indeed, in all but one case, growth rates were positive, in contrast to wildtype parasites (**Supplementary Fig. S3**). Furthermore, growth rates exhibited dose-dependency as populations of parasites that had been previously exposed to higher doses of atovaquone grew better, and at the highest doses approached rates characteristic of wildtype parasites in non-drug treated hosts (experiment 1 (log10 transformed drug dose): F_1,13_ = 14.5, p = 0.002; experiment 2 (log10 transformed drug dose): F_1,58_ = 74.6, p < 0.001, **Fig. 3B**). Thus, higher drug doses led to parasite populations that had higher levels of resistance.

The effects of dose on resistance emergence were not obviously related to parasite kinetics during drug treatment. Drug treatment with any dose of atovaquone resulted in significant decline of parasite numbers, reducing populations by nearly an order of magnitude per day (**Supplementary Fig. S4a**). Parasites were cleared more rapidly in experiment 1 (C57BL/6 mice) than they were in experiment 2 (Swiss Webster mice; F_1,82_= 24.3, p < 0.001). Rates of parasite clearance in drug-treated mice were unaffected by dose in experiment 1 (F_1, 35_ = 0.95, p = 0.3) and, unexpectedly, decreased with increasing dose in experiment 2 (Swiss Webster mice; F_1,76_ = 11.3, p = 0.001). Population sizes at the start of treatment differed significantly between our two experiments (F_1,101_ = 26.4, p < 0.001, **Supplementary Fig. S4b**).

Drug treatment had no effect on the survival of C57BL/6 mice but rescued Swiss Webster mice (**Supplementary Fig. S5**). Among treated mice, dose did not affect survival (p = 0.4, p = 0.6 in experiment 1 and 2, respectively), but it did affect anemia. In the more resilient mouse strain (C57BL/6), higher drug doses limited anemia during the acute stages of infection (F_1,33_ = 14.1, p = 0.001, **Supplementary Fig. S4c**), but not when infections relapsed (F_1,18_ = 0.22, p = 0.64, **Supplementary Fig. S4d**). In the less resilient Swiss Webster mice, dose had no impact on acute-phase anemia (F_1,68_ = 0.1, p = 0.8, **Supplementary Fig. S4c**), but higher dose proved to be more protective when infections relapsed (F_1,64_ =4.1, p = 0.05, **Supplementary Fig. S4d**). In both experiments, relapse led to significantly more anemia in chronic stage infection compared to non-relapsed mice (experiment 1: F_1,35_ = 12.8, p = 0.001; experiment 2: F_1,76_ = 11.5, p = 0.001), although these differences were more noticeable in Swiss Webster mice.

### Timing of atovaquone treatment and parasite population size

In our second set of experiments (experiments 3-5), we held atovaquone dose constant and manipulated timing of treatment or inoculum size to assess the effect of timing and population size on infection relapse and resistance evolution (summarized in **Fig. 1C-E, Supplementary Table S1, Fig. 4**).

**Fig 4.**
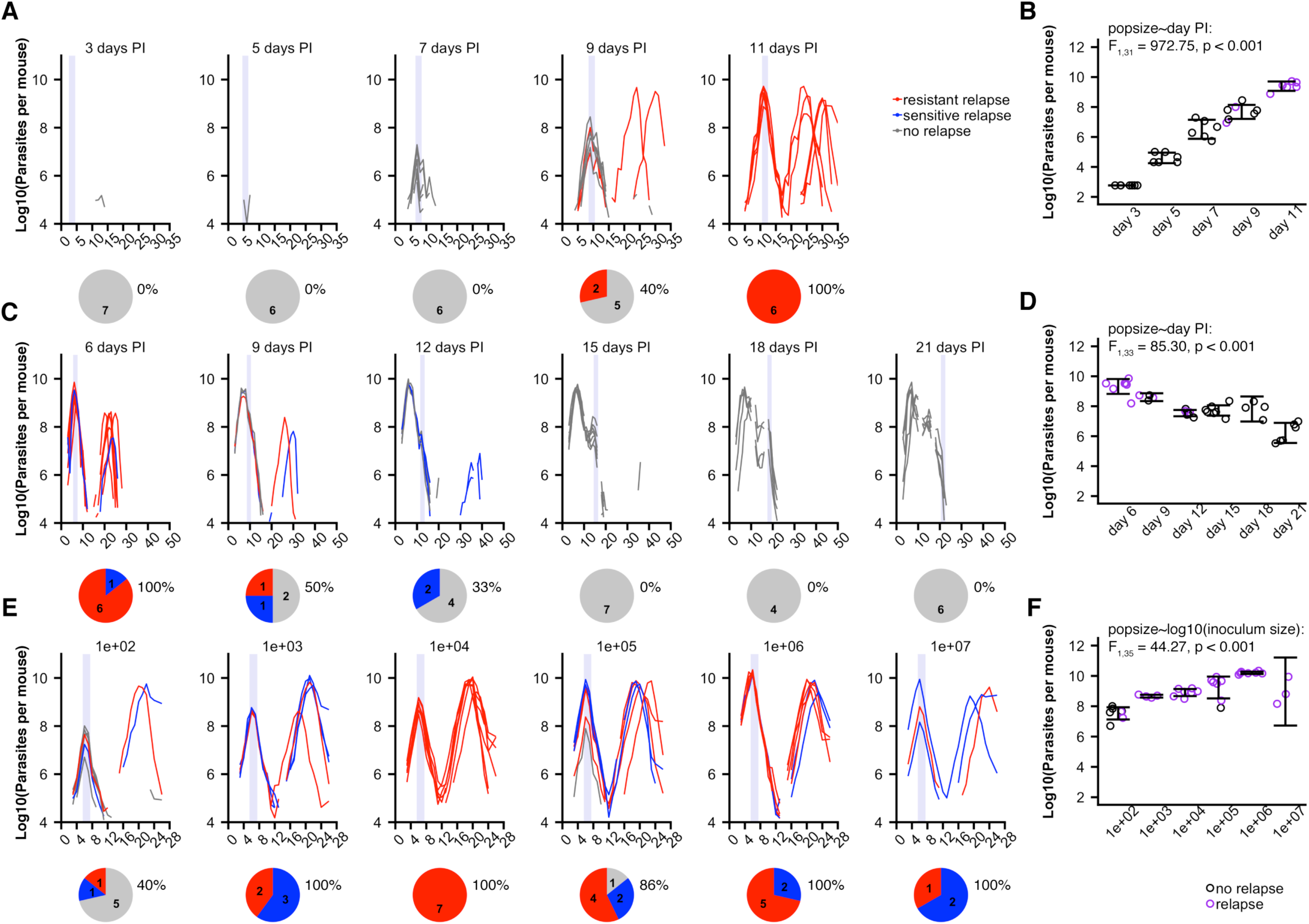
Individual parasite dynamics for experiments 3-5 and treatment failure and resistance. **Left panels,** parasite dynamics for all individual mice in experiments 3 **(A)**, 4 **(B)** and 5 **(C)**. Colors indicate the consensus sequence of the Qo2 region of the *cytb* gene (wildtype, blue; resistance-associated mutants, red; grey, non-relapsing infections). Pie diagrams show proportion and total numbers of mice in each category. Percentages show proportion of total infections which relapsed per treatment. Light purple bars, timing of drug treatment. **Right panels**, population sizes at time of drug treatment for experiment 3 **(D)**, 4 **(E)** and 5 **(F)** for each treatment; purple, infection relapsed; black, no relapse. Error bars represent 95% confidence intervals, boxes represent averages for a given experimental group. Statistics demonstrate whether population size increased with time since inoculation **(D &E)** or with inoculum size **(F)**.

In experiment 3, Swiss Webster mice were infected with 100 parasites and treated at various times before peak parasitemia (**Fig. 1C, Table 1**). Resistance only emerged in later-treated infections (resistance∼experimental group: χ_4,31_^2^ = 27.61, p < 0.0001, **Fig. 4A**). These infections had larger population sizes when treatment started (population size∼day post-infection: F_1,31_ = 972.75, p < 0.001, **Fig. 4B**). Within experiment groups, there was variation in population size at time of treatment, but this variation did not additionally predict resistance emergence (χ_1,31_^2^ = 0.52, p = 0.47).

In experiment 4, C57BL/6 mice were infected with a million parasites and treated at various points after peak infection (**Fig. 1D, Table 1**). Treatment failure and resistance emergence was less likely in later-treated infections (relapse∼experimental group: χ_5,33_^2^ = 29.62, p < 0.001; resistance∼experimental group: χ _5,33_^2^ = 24.33, p < 0.001, **Fig. 4C**). Population sizes at the start of drug treatment were also smaller in the later-treated infections (population size∼day post-infection: F_1,33_ = 85.30, p < 0.001, **Fig. 4D**). Within experiment groups, there was variation in population size at time of treatment, but this variation did not additionally predict the likelihood of relapse or resistance emergence (relapse: χ_1,33_^2^ = 1.27, p = 0.3, resistance: χ_1,33_^2^ = 0.03, p = 0.9).

In experiment 5, Swiss Webster mice were inoculated with different numbers of parasites to generate different populations sizes when drug treated on days six and seven post-infection (**Fig. 1E, Table 1**). The likelihood of relapse and resistance differed among experimental groups, with resistance most likely at intermediate inoculum sizes (relapse∼experimental group: χ_5,35_^2^ = 18.3, p = 0.003; resistance∼experimental group: χ_5,35_^2^ = 15.2, p = 0.01, **Fig. 4E)**. Here, population sizes at time of treatment did not continuously increase as inoculum sizes increased, reflected by a significant quadratic term in the relationship (population size∼log10(inoculum size)^2^: F_1,34_ = 10.0, p = 0.003, **Fig. 4F**). Within experiment groups, there was variation in population size at time of treatment, but this variation did not additionally predict the likelihood of relapse or resistance emergence (relapse: χ_1,35_^2^ = 1.96, p = 0.16, resistance: χ_1,35_^2^ = 0.01, p = 0.9). Note that the variation in population size at time of treatment was substantially less in this experiment than in experiments 3 and 4 (σ^2^ = 0.92 versus σ^2^ = 1.17 for experiment 4 and σ^2^ = 5.81 for experiment 3, generalized least squares model allowing for variance structure, **Supplementary Fig. S6**).

Thus, in all three experiments, relapse, resistance emergence and population size at the start of atovaquone treatment differed among experimental groups. We therefore asked if population size per se impacted relapse and resistance. Across all experiments, relapsing infections were characterized by larger population sizes at time of treatment (experiment 3: F_1,31_ = 25.77, p << 0.0001; experiment 4: F_1,33_ = 24.01, p << 0.0001; experiment 5: F_1,35_ = 20.17, p << 0.0001, **Fig. 5**). Population size, however, did not determine whether those relapses were dominated by resistant or sensitive parasites (experiment 4: F_1,10_ = 3.90, p = 0.08; experiment 5: F_1,29_ = 0.04, p = 0.83, **Fig. 5**) but did increase the risk of resistance generally (**Supplementary Fig. S7**).

**Fig 5.**
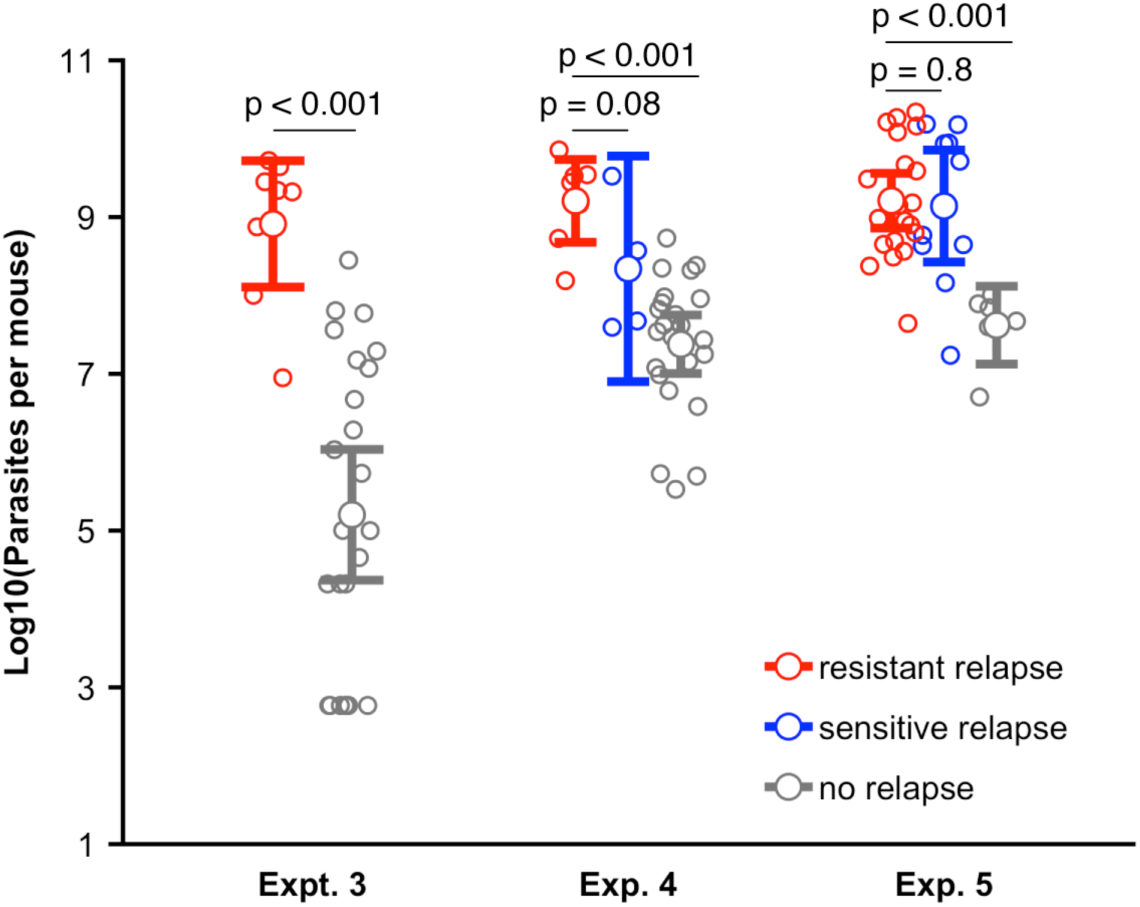
Treatment outcome as a function of parasite population size at start of atovaquone treatment. Pathogen population sizes for non-relapsing (gray) infections, and for relapsing infections that were genotyped as drug-resistant (red) and or wildtype/drug-sensitive (blue). In all three experiments, infections that relapsed had significantly larger parasite population sizes when treatment started, but resistant relapses were no more likely to emerge from larger populations than were sensitive relapses. Error bars are 95% confidence intervals.

### Resistant mutants

In all cases, non-wildtype genotypes from relapsed populations contained mutations previously reported to result in high level resistance to atovaquone in *in vivo* experiments or from human field data [51,61–63]. All detected mutations were confined to a short region of the Qo2 domain spanning only six amino acids in length (**Supplementary Fig. S1**). Various SNPs led to seven non-synonymous mutations (F267I/V, Y268C/N/S, L271V and K272R) resulting in a total of ten different consensus sequences across all five experiments (**Supplementary Table S2**). In most cases, consensus sequences reflected the presence of a single amino acid change, but in some instances, two mutations were found to co-occur in the consensus sequence. This was most often the case with mutations L271V and K272R, the other example being a single infection with mutations Y268C and L271V in experiment 2 (**Supplementary Table S2**). These are likely double mutant haplotypes because the mutations appear to be equally abundant as judged from the electropherogram, but we cannot exclude the possibility that these are mixtures of two populations each with single mutation haplotypes. Our data also suggested that in some cases there was parasite population diversity within single mice (**Supplementary Text**).

A large diversity of mutations were selected even within the context of a single dose across different mice. This is reflected in the proportion of mice which harbored parasites of various genotypes at different doses for experiments 1 and 2 (**Supplementary Figure S8a**). While wildtype genotypes dominated relapsing populations at low doses, higher doses resulted in a greater proportion of mice harboring parasites of the Y268C genotype (drug dose: χ_1,5_^2^ = 22.24, p < 0.001). Moderate doses (8 mg/kg) resulted in the largest number of represented consensus genotypes within experiments 1 and 2, with a total of seven represented genotypes.

## DISCUSSION

Using an experimental system where *de novo* resistance readily emerges *in vivo*, we found that while higher doses of atovaquone led to an increased risk of resistance in only one of two experiments, higher level resistance (as reflected by our phenotyping data) was increased in both. Timing of treatment also impacted resistance emergence, but we were unable to demonstrate that this was a simple function of population size. We take each of these results in turn.

### Dose and the emergence of resistance

When we varied atovaquone concentration (experiments 1 and 2), increasing dose led to the evolution of increasingly higher-level resistance, as measured by the growth rates of relapsing parasites in drug-treated tester mice (**Fig. 3B**). Presumably high-level resistance emerges only at the highest drug doses because mutations conferring the greatest levels of resistance have accompanying costs that put parasites at a competitive disadvantage in the absence of strong drug pressure, meaning that these mutations can only emerge when parasites with mutations that confer lower-level resistance are killed. Indeed, resistance is accompanied by documented fitness costs in this system, which differ between haplotypes of mutants [57], likely arising due to differences in reduced binding affinity of atovaquone for parasite *cytb* [67].

At the genetic level, the proportion of infections that harbored the Y268C mutant also increased with dose (**Supplementary Fig. S8a**). This mirrors the phenotypic resistance results as the substitution of a tyrosine for a cysteine at position 268 is associated with one of the highest levels of resistance to atovaquone in both *in vivo* and *in vitro* studies [51,63,67]. However, the Y268C mutation was not the sole cause of the high-level resistance that emerged at higher doses. Other mutations were recovered from relapsing infections in mice treated with high doses, but were less common (**Supplementary Fig. S8a**). Furthermore, it is possible that the high-level resistance in some infections was conferred by sub-populations with alternative or additional mutations each of which existed at frequencies insufficient to feature in our consensus sequence of the Qo2 domain generated by Sanger sequencing. Indeed, we frequently detected evidence of mixed infections when we examined sequencing electropherograms, particularly in infections where high-level resistance emerged (**Supplementary Fig. S8b**). It is also possible that some high-level resistance (**Fig. 3B**) was conferred by mutations outside of the Qo2 domain. Mutations elsewhere in *cytb* are known to be associated with atovaquone resistance, though none with high-level resistance [57]. Full and deep genome sequencing of the parasites that emerged in our experiments could be revealing but accounting for the entire genetic basis of evolved atovaquone resistance was not an aim of our study.

In experiments 1 and 2, dose had contrasting impacts on the probability that parasites mutant at the Qo2 domain would emerge (**Fig. 2, 3A**). Dose had no impact in our first experiment, but in our second, the likelihood of mutations arising in the Qo2 domain increased with dose. There are several possible explanations for these experiment differences, and while we did not explicitly test for it, differences in mouse strain seem most likely. It is well known that antimicrobials are less efficacious and resistance evolution is more likely in immune-impaired individuals [68–70], and indeed, parasite populations were smaller at the time of treatment and drug-induced clearance rates were greater in the more resilient C57BL/6 mice (experiment 1) than in Swiss Websters (experiment 2). Differences in the relative contribution of genetic drift and selection following drug treatment or in the relationship between dose and resistance may account for this. Going forward, host heterogeneity is something to consider in experimental analyses of resistance-retarding treatment regimens.

Our results differ from many *in vitro* studies and a few *in vivo* studies in which the evolution of resistance was reduced at higher doses [71–73]. The discordance between these studies and ours likely arises from the existence of the so-called inverted U, where resistance evolution is minimized at very low or very high doses and maximized somewhere in-between [3,4]. Presumably, if we had increased concentrations still further, we would have reached a point where there were no mutants capable of surviving and so no resistance would have emerged despite the intense selection. However, with our formulation, we were up against the upper bound of the therapeutic window, the point at which the drug itself becomes harmful. Atovaquone has limited solubility (even in DMSO) and the solvent itself is toxic [74], particularly in mice already suffering from malaria infection (data not shown). It is possible that higher doses, perhaps delivered via oral administration, would reveal the full inverted U. Nonetheless, our data caution against the simple mantra that more is better. Just because the inverted U is well understood in theory and well verified *in vitro*, biological realities can make it challenging to implement [4]. The right dose need not be the highest dose [3,4,75].

### Timing and the emergence of resistance

When the timing of treatment was altered, infections treated closest to the time of peak parasite density were more likely to result in resistance (here treated only as binary genotypic resistance). As such, treating early or very late in infection resulted in decreased probabilities of resistance emergence (experiment 3 and 4, respectively; **Fig. 4A and C**). The simplest explanation for the effect of timing on resistance evolution is the size of the pathogen population at the time of treatment: population sizes are small early or late in infection (**Fig. 4B and D**), consistent with the general principle that larger population sizes are more likely to contain drug-resistant mutations on which selection can act [40–43]. However, for infections treated after peak densities (experiment 4), an alternative explanation is that developing immune responses decrease the opportunity for resistance emergence. Models of pathogen growth in rodent hosts have shown that pathogen dynamics are disproportionately regulated by resource limitation (e.g. red blood cells) early in infections, but after 10 days immune regulation dominates [76]. Relapses may thus be substantially restricted by active immunity when treatment is given later. Post-treatment relapses were indeed smaller when treatment began on day 12 post infection than those following treatment on days 6 and 9 post infection (**Fig. 4C**).

Even if immunity restricts resistance emergence later in infections, our data suggests that when the parasite population is expanding, the earlier treatment occurs, the better (experiment 3, **Fig. 4A**). The most likely explanation for this is the smaller populations at the time of treatment. We directly tested this by altering the initial inoculum size to generate different populations on a fixed day of treatment (experiment 5). We found that resistance evolution was maximized at intermediate inoculum sizes (10 000 parasites, **Fig. 4E**). While we expected that increasing inoculum sizes would result in concurrent increases in parasite population size at the time of drug treatment, they were similarly maximized at intermediate inoculum sizes, with the largest population sizes generated by an inoculum size of 10^6^ (**Fig. 4F**). The reason for this is unclear, and may reflect some biases in our experimental set-up. Indeed, only three of eight mice survived the acute-stage of infection in our highest inoculum size manipulation (10^7^). As such, the mice which we included in our analysis may reflect mice that received a lower inoculation by chance or instances in which mice were able to better control their parasite populations. Despite this, our data suggest that the risk of resistance was more related to population size than inoculum size. However, resistance risk did not differ between mice given an inoculum size of 10 000 or 10^6^ parasites, irrespective of significant population size differences (population size∼experimental group, F_1,13_^2^ = 169.55, p < 0.001, **Fig. 4E**; resistance∼experimental group, F_1,13_^2^ = 2.4, p = 0.15, **Fig. 4E)**.

Taken together, this suggests that the relationship between population size and resistance is not altogether straightforward, or that its effect may be overwhelmed by other factors when differences in population sizes are small or moderate. We found a correlation between resistance emergence and population size in each of our three experiments (**Supplementary Fig. S7**), but we cannot statistically disentangle this from the effects of our experimental treatments *per se* (**Supplementary Table S1**). This might be a detection issue. Manipulating inocula sizes did not generate populations as small as those in the other experiments (**Fig. 4B,D,F; Supplementary Fig. S6**), and that more limited between-group variation may have been insufficient to reveal a clear relationship between population size and resistance risk. Likewise, in all three experiments, the within group variation in population size was small relative the between group variation (**Fig. 5**), again perhaps limiting our ability to detect an effect. Further experiments are required to resolve this. In the meantime, all we can say is we do not have direct experimental evidence that treating early prevents resistance emergence because populations are small, even though that seems the most obvious explanation.

## ‘RIGHT’ DOSE, ‘RIGHT’ TIMING?

Our experiments are some of the very few that directly measure *de novo* resistance emergence *in vivo* as a function of treatment regimen. What do they say about the ‘right’ dose and ‘right’ timing of patient treatment? Our aim was to advance the science of regimen choice, and we cannot expect to identify particular regimens that will be optimal in real-world settings: details matter [4]. Nevertheless, we can ask for our animal model, what dose and timing best manage resistance emergence while simultaneously maximizing host health, and we can consider the implications were those findings to generalize.

We found unequivocal evidence that the timing of treatment matters (**Fig. 4A and B**), so that from the perspective of resistance management within a host it is better to treat as early as possible or to withhold treatment until immunity has built up. This accords with theory [34,36,77] and the long standing ‘hit-early’ orthodoxy [78]. The practicalities of hitting early (or late) will depend on the circumstances. It may be that once symptoms are detected, infections are already advanced to the point where resistance emergence is a real risk. The feasibility of withholding treatment until immunity is building will be very situation specific.

The ‘right’ dose differed between experiments. In our two experiments where we varied drug concentration, atovaquone dose affected resistance emergence (**Fig. 2 and 3**) and, above the very lowest doses, had negligible impact on host health (**Supplementary Fig. S4c and d**). In our first experiment, however, increasing drug dose resulted in fewer treatment failures overall (**Fig. 2A**), and as relapses resulted in greater anemia later in infection, there is an argument that the highest doses were the ‘right’ dose, despite the consequences for resistance. In our second experiment where increasing dose did not decrease the probability of treatment failure, it would seem the best option is to use a low dose to allow the possibility of further treatment during relapse. Alternatively, an intermediate low dose (i.e. 4 mg/kg) could be selected if patient health was a more important consideration as doses above 2 mg/kg provided some protection against anemia during relapse (**Supplementary Fig. S4d**). Assessing whether this slight clinical gain (mice were still significantly anemic) is worth the additional risk of resistance is an example of choices that arise due to different effects of dose on resistance risk and clinical health. These contrasting conclusions arising in even our highly simplified setting demonstrate that identifying the “right’ dose is a non-trivial problem.

Minimally, then, our data demonstrate the need for continued evaluation of what constitutes ‘appropriate’ antimicrobial treatment [3,4,32,33,77]. Many questions remain. For example, the impact of host immunity on pathogen dynamics during drug treatment is a big knowledge gap [33,79]. This is especially important as current attempts to search for new antimicrobials continues to focus on fast killing compounds [80] and drug development pipelines make little room for testing evolution of resistance in *in vivo* scenarios.

## ACKNOWLEDGEMENTS

We thank members of the Read group for discussion, especially E. Hansen, D. Kennedy, L. Pollitt and N. Wale.

