## Supplementary Text for "The effect of drug dose and timing of treatment on the emergence of drug resistance *in vivo* in a malaria model"

***Sequencing relapsed infections and quantifying mixed infections (experiment 1 and 2)***

Parasite DNA from samples at peak relapse were amplified using forward: 5’ TCC TTT AGG GTA TGA TAC AGC 3’ and reverse primers: 5’ CTT GAG GTA ATT GAC ATC CTA TC 3’, at a final concentration of 0.3 μM using the Qiagen *Taq* PCR Core Kit and purified via the Qiagen QIAquick PCR Purification Kit prior to sequencing. To confirm that the initial parasite populations from which we selected resistance were indeed wildtype at the Qo2 region, we amplified and sequenced parasite DNA from twenty infections in experiments 1 and 2 on the first day of drug treatment. Only the wildtype genotype was detected. We also sequenced parasites from untreated infections later in infection (ten mice in experiment 1 on day 13 post-infection and a single mouse in experiment 2 on day 20 post-infection) and detected only the wildtype Qo2 genotype. Thus, in the absence of drug treatment, any mutations in that region were at densities below our detection threshold.

In almost all cases sequencing of the amplicon in both the forward and reverse direction resulted in identical sequences, however, a minority of samples differed with respect to either the majority genotype represented or in the presence of mixed infections detected. In all main text analyses, we refer to the majority genotype, obtained from Sanger sequencing in the forward direction. In our analyses here, we consider the presence of mixed infections, defined as: infections in which forward and reverse sequences differed and/or and minority peaks were detected in sequencing (at least 25% peak similarity, as detected by Geneious® version 9.1.8 and confirmed via manual inspection).

Mixed infections were common within our relapsing populations. We re-classified sequences from infections into three categories: wildtype (no detection of minority peak alleles), mutant (no detection of minority peak alleles) and mixed (evidence of minority peak alleles); based on both the forward and reverse sequences. Data from our phenotypic measures of resistance suggested that phenotypic resistance varied depending on genotype in experiments 1 and 2 (genotype: F_9,57_ = 7.32, p < 0.001, **Supplementary Fig. S8b**). We found that phenotypic resistance was determined in part by the presence or absence of mixed infections. Wildtype only infections were found to have the lowest slopes of parasite growth in the presence of drug in naïve mice (0.02 +/- 0.17, 95% confidence interval), followed by mixed infections (0.58 +/- 0.08, 95% confidence interval) and finally mutant only infections (0.63 +/- 0.10, 95% confidence interval).

The large spread in phenotypic variance for wildtype genotypes is likely because some of these relapsing populations contained sub-dominant clones. Given the low ability of Sanger sequencing to fully resolve mixed infections, it is likely that our estimations of mixed infections are highly conservative and that drug treatment resulted in relapse with highly diverse pathogen populations that differed in genotype and relative frequency in most cases. Selection here thus occurs as a soft selection sweep involving multiple and independent origins of the same or related alleles that confer drug resistance. Soft sweeps have been theorized to be common under scenarios of high mutation rates and/or high selection coefficients^1,2^ and have even been predicted to occur given the complex biology of mitochondrially encoded atovaquone resistance^3^.
