## Supplementary Table S2 for "The effect of drug dose and timing of treatment on the emergence of drug resistance *in vivo* in a malaria model"

**Supplementary Table S2. Consensus haplotypes at the Qo2 domain of the *cytb* gene during relapses**. Tabulated values are the number of infections. Underlined amino acids in the first column denote mutated amino acids as compared to the wildtype sequence.

| **Haplotype** | **Mutations** | **Previously reported*** | **Total Numbers** | **Total Numbers** |
| --- | --- | --- | --- | --- |
|  |  |  | ***Experiment 1 & 2*** | ***Experiment 3-5*** |
| FYAMLK (wildtype) | - | - | 29 | 12 |
| FCAMLK | Y268C | yes^a,c,d,e,f^ | 29 | 5 |
| FYAMVR | L271V + K272R | yes^a,c,e,f^ | 18 | 12 |
| FNAMLK | Y268N | yes^b,c,e,f^ | 7 | 10 |
| FSAMLK | Y268S | yes^d,f,g^ | 5 | - |
| FYAMVK | L271V | yes^t^ | 5 | 3 |
| IYAMLK | F267I | Yes^a^ | 4 | - |
| FYAMLR | K272R | yes^t^ | 3 | 3 |
| FCAMVK | Y268C + L271V | yes^t^ | 1 | - |
| VYAMLK | F267V | no | 1 | 2 |

**^*^** column indicates study in which mutations were reported: ^a^ Srivastava *et al* 1999 (*P. yoelii)* , ^b^ Afonso *et al* 2010 (*P. chabaudi*), ^c^ Siregar *et al* 2008 (*P. bergheii*), ^d^ Musset *et al* 2006, ^e^ Nuralitha *et al* 2017, ^f^ Nuralitha *et al* 2016, ^g^ Korsinczky *et al* 2000. ^t^ denotes mutations where mutation has been previously reported, but not in the specific haplotype indicated here.
