## Supplementary Fig S3 for "The effect of drug dose and timing of treatment on the emergence of drug resistance *in vivo* in a malaria model"

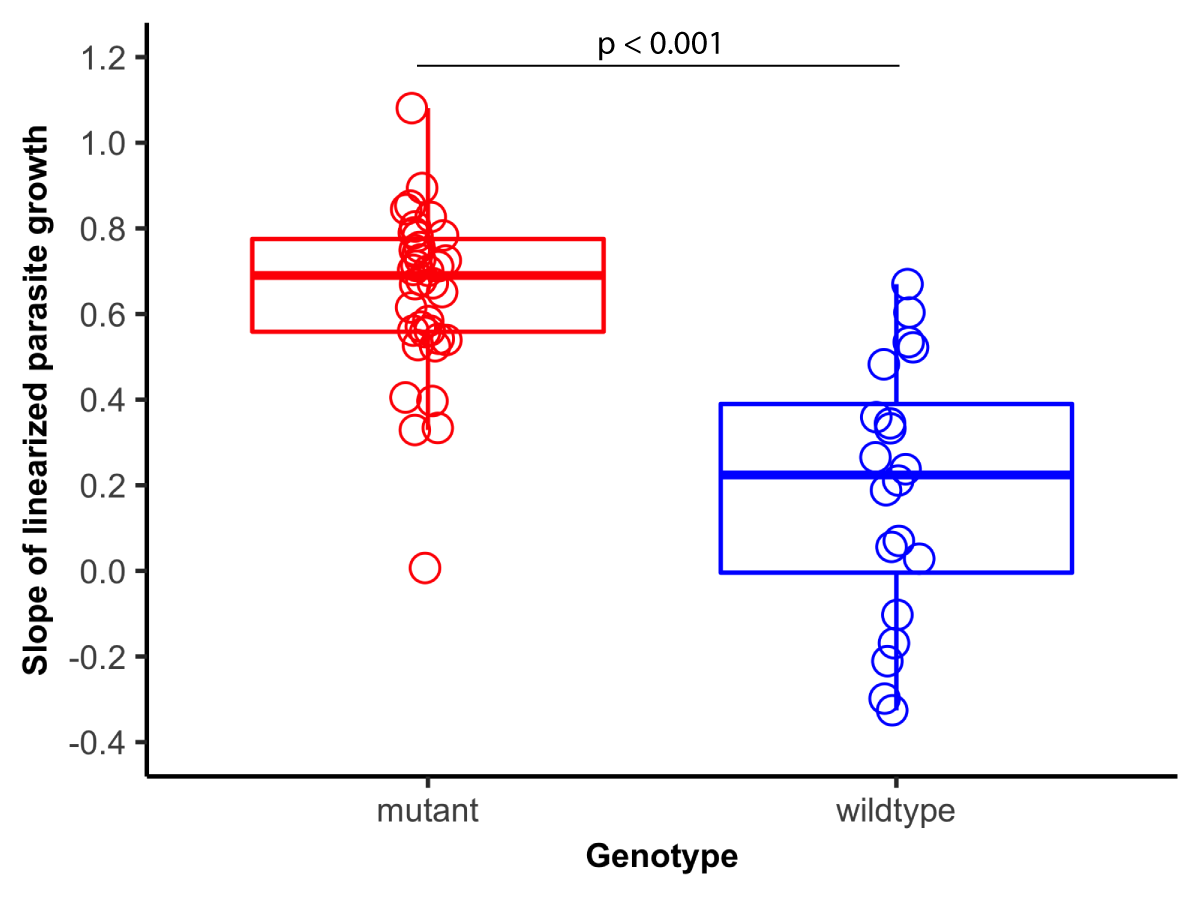

**Supplementary Fig S3. Growth rates of Qo2 mutant** (red) **and wildtype** (blue) **parasites in naïve drug-treated mice**.
