## Supplementary Fig S8 for "The effect of drug dose and timing of treatment on the emergence of drug resistance *in vivo* in a malaria model"

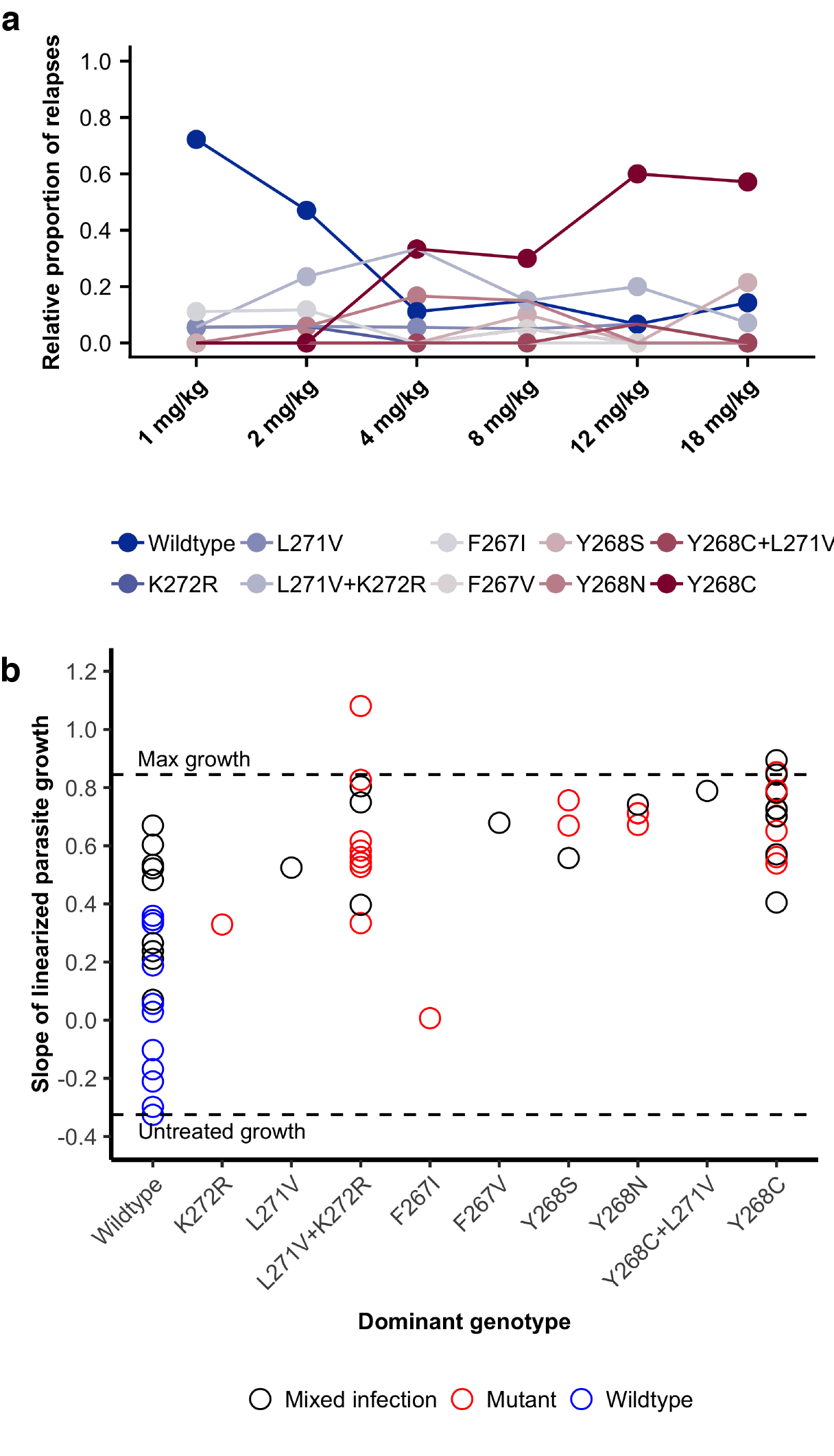

**Supplementary Fig S8. High-level resistance is associated with a diversity of mutations.** (**a**) The proportion of relapses due to a specific genotype are shown across dose for data combined between experiments 1 and 2. (**b**) Parasite growth of relapsed infections in atovaquone treated tester mice is shown for experiments 1 and 2 as a function of their genotype. Colors show whether or not evidence of mixed infections were found for each individual mouse relapse (black indicating mixed infection). Blue is wildtype drug-sensitive with no evidence of mixed infection, red is mutant drug-resistant with no evidence of mixed infection.
