## Supplementary Fig S7 for "The effect of drug dose and timing of treatment on the emergence of drug resistance *in vivo* in a malaria model"

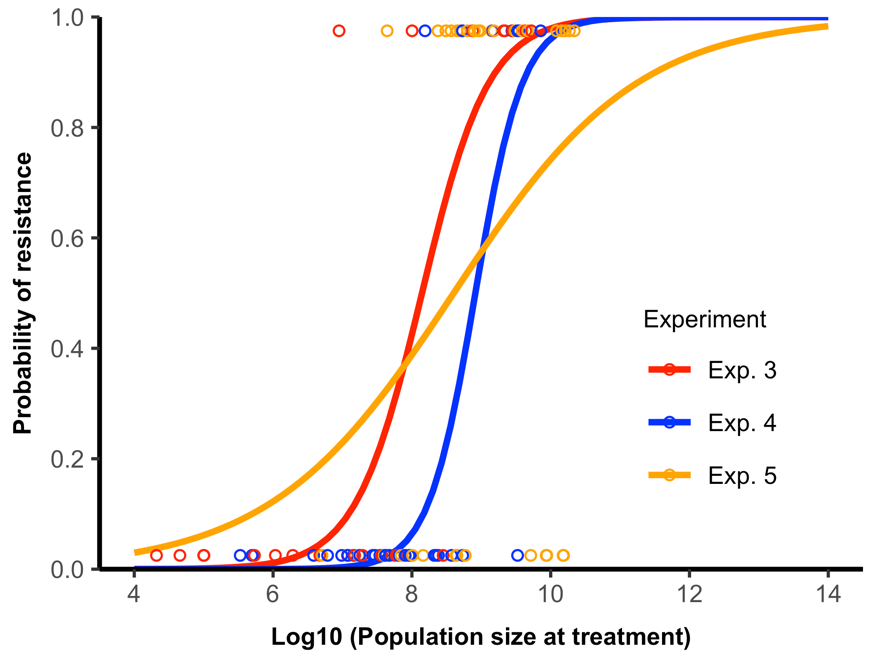

**Supplementary Fig S7. Population size at time of treatment and resistance emergence.** Parasite population size at the time of treatment plotted against the probability of resistance emergence across all experimental groups in experiments 3 (red), 4 (blue) and 5 (yellow). Fitted curves reflect GLM model predictions fitted for each experiment.
