## Supplementary Fig S6 for "The effect of drug dose and timing of treatment on the emergence of drug resistance *in vivo* in a malaria model"

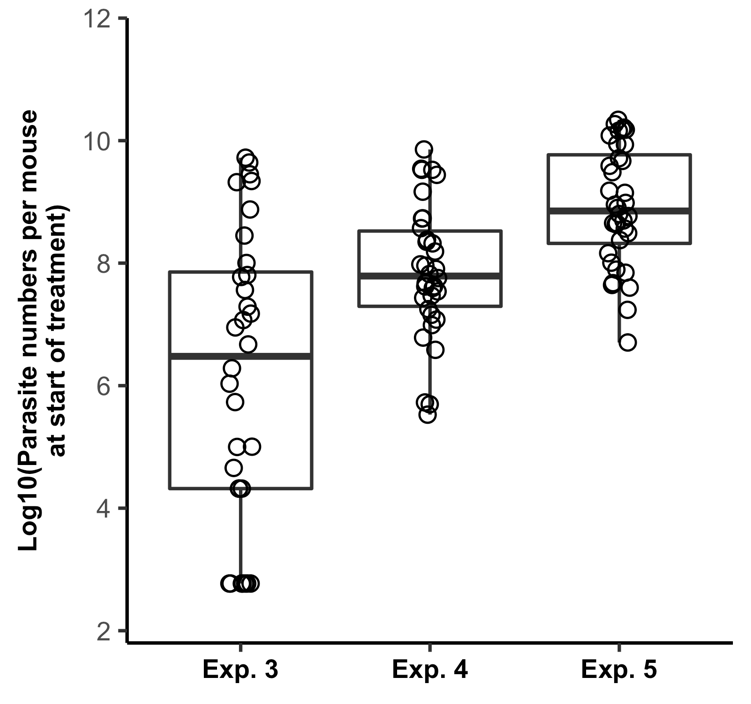

**Supplementary Fig S6.** **Parasites per mouse at start of atovaquone treatment for experiments 3-5**. Horizontal bold lines show the mean, and boxplots show variance of population sizes generated by each experimental manipulation. σ^2^ = 5.81 for experiment 3, σ^2^ = 1.17 for experiment 4 and σ^2^ = 0.92 for experiment 5 via generalized least squares model allowing for variance structure.
