## Supplementary Fig S5 for "The effect of drug dose and timing of treatment on the emergence of drug resistance *in vivo* in a malaria model"

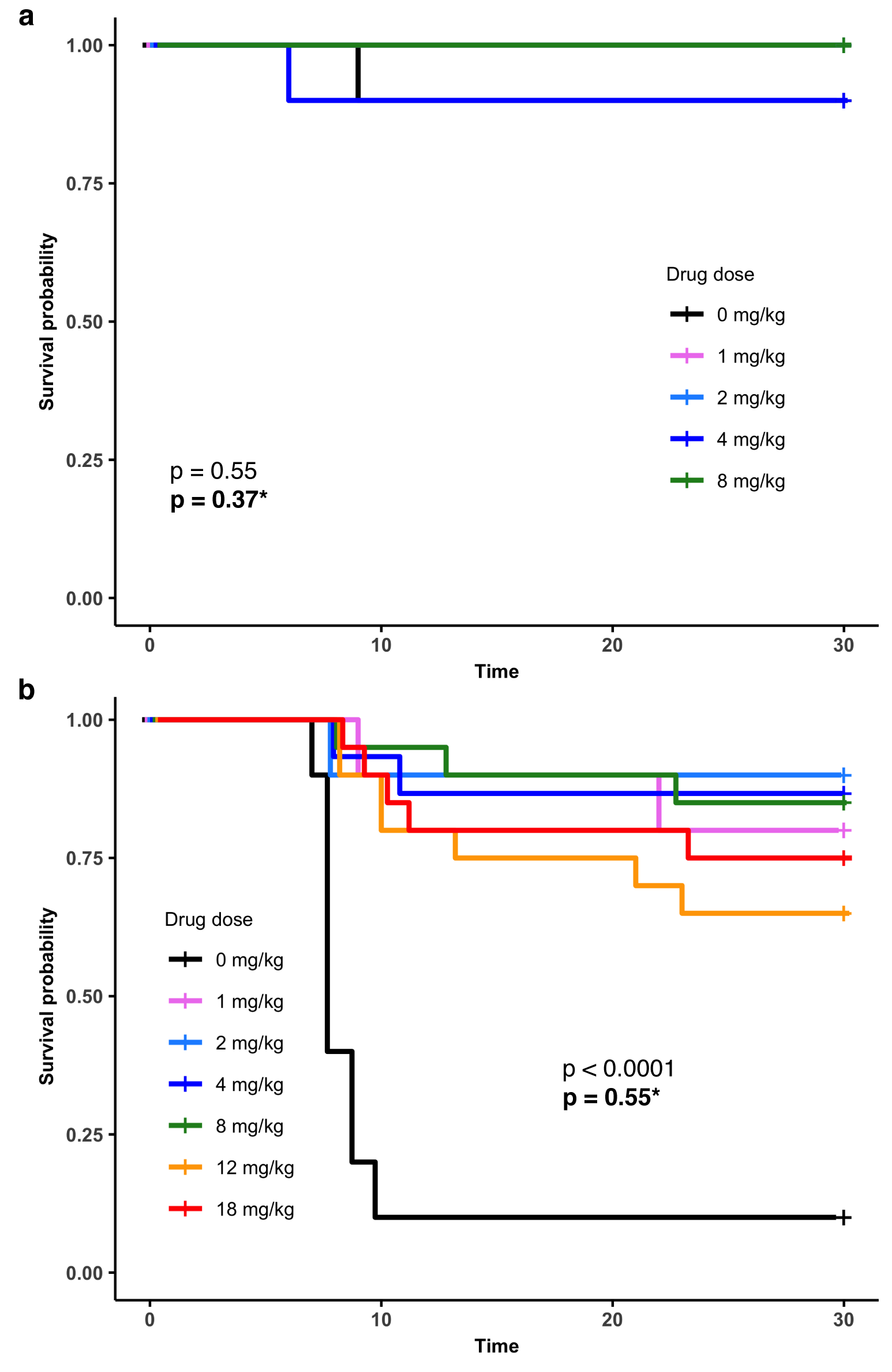

**Supplementary Fig S5.** **Kaplan-Meier survival curves for experiments 1 and 2**. Survival curves are shown for mice in experiment 1 (**a**) and 2 (**b**). Unbolded p-values represent the effect of all experimental groups on survival probability; *Bold p-values when omitting no-drug controls.
