## Supplementary Fig S4 for "The effect of drug dose and timing of treatment on the emergence of drug resistance *in vivo* in a malaria model"

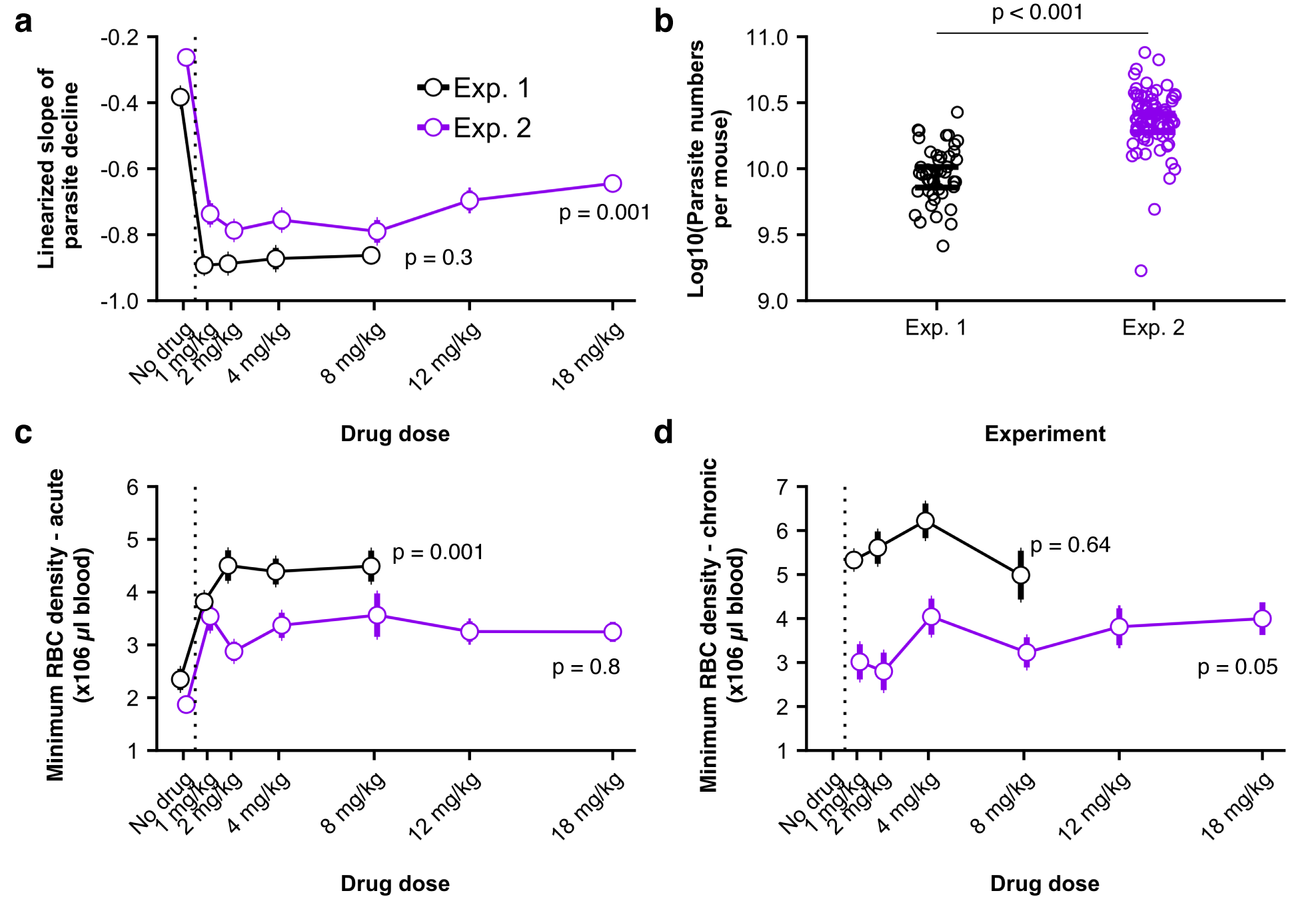

**Supplementary Fig S4. Effects of atovaquone treatment on parasite dynamics and host health**. Effects of treatment or experiment are shown for experiments 1 (black) and 2 (purple) in each panel. **(a)** Rate of parasite clearance during and immediately after drug treatment (days 6-10 post-infection). **(b)** Average population sizes at time of treatment**.** (**c,d**) Minimum red blood cell densities for individual mice during acute (**b**) and chronic (**c**) stages of infection. Only data from relapsed mice are shown for the chronic stage of infection. Dotted black lines separate drug treated and control untreated mice. Error bars denote 95% confidence intervals.
