## Supplementary Fig S2 for "The effect of drug dose and timing of treatment on the emergence of drug resistance *in vivo* in a malaria model"

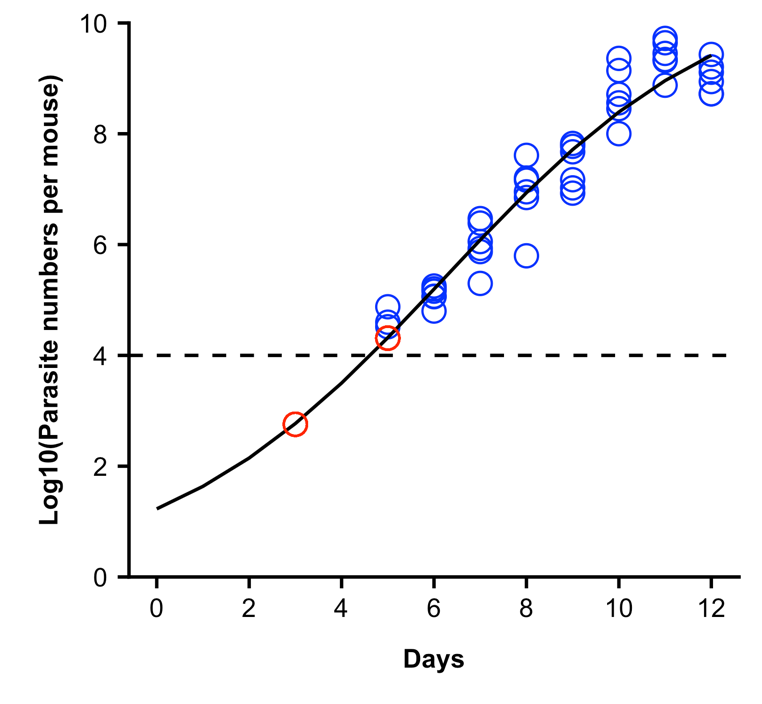

**Supplementary Fig S2. Estimating population sizes at time of treatment for early treatment groups in experiment 3**. We fit a 3-parameter logistic model to data from untreated mice during parasite growth following inoculation of an estimated 100 parasites. In blue is data from our untreated mice, while red points are estimates for parasite numbers on days when parasites were below the level of detection in our qPCR assay (dotted line), as predicted by model fit.
