## Supplementary Fig S1 for "The effect of drug dose and timing of treatment on the emergence of drug resistance *in vivo* in a malaria model"

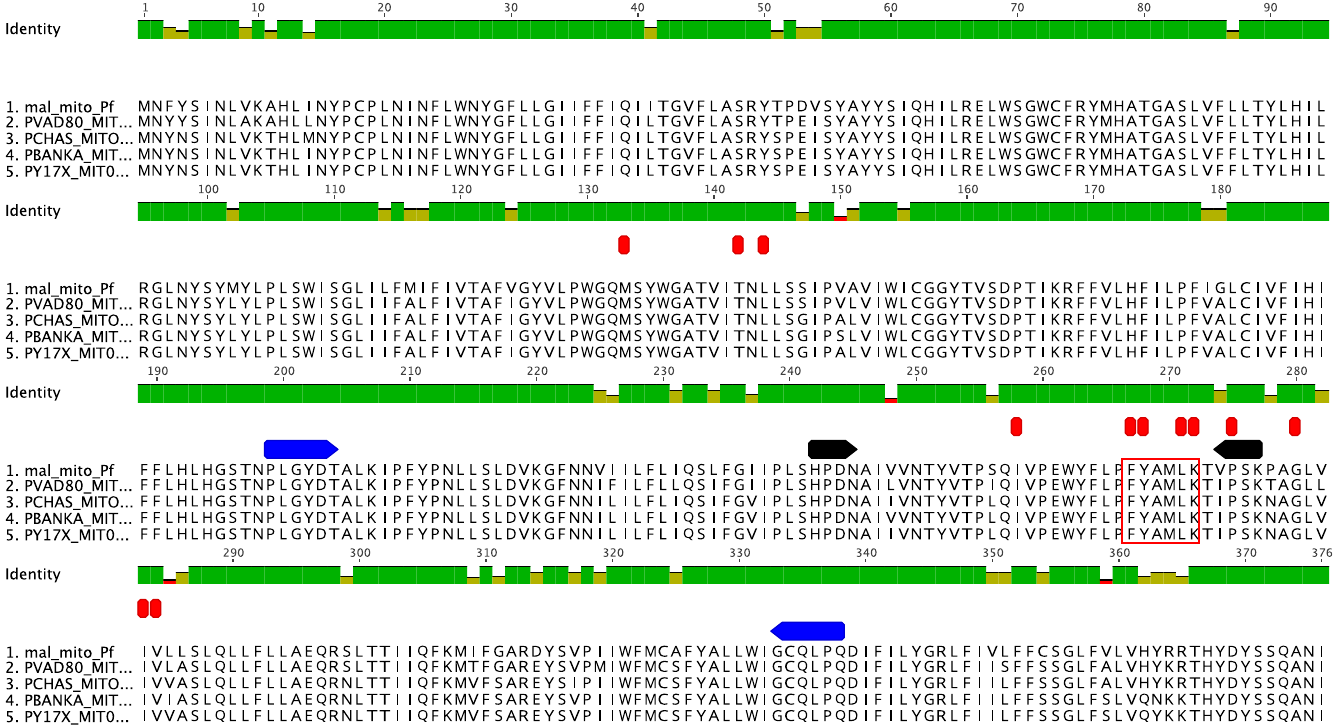

**Supplementary Fig S1. Five species alignment of the *cytochrome b* gene from malaria parasites.** Sequences of the full-length *cytochrome b* gene are shown for a representative atovaquone sensitive (wildtype) strain of *P. falciparum* and *P. vivax* (human infecting species; PlasmoDB reference IDs: Pf_M7661101900.1, PVAD80_MIT0003.1, respectively) and *P. yoelli, P. chabaudi* and *P. berghei* (rodent infecting species; PlasmoDB reference IDs: PYYM_MIT00900.1, PCHAS_MIT01800.1, PBANKA_MIT01900.1, respectively)*.* Sequences were aligned via Geneious version 9.1.8. Sequence homology is represented above the alignment with green bars, indicating large levels of conservation even among these distantly related species. Blue arrows indicate positions of our forward and reverse primers used in sequence amplification and sequencing. Red bars indicate known and reported mutations associated with resistance to atovaquone in previously reported *in vitro* and *in vivo* studies. Black arrows indicate the quinone binding 2 (Qo2) region of the gene in which high level resistant mutations have been reported. The red box indicates where all the presently reported mutations were located.
